# Haloperidol-induced impairment on working memory capacity affecting long term memory performance: the binding hypothesis

**DOI:** 10.1101/2019.12.24.888073

**Authors:** Regina V. Guarnieri, Marcus Vinicius Alves, Ana Maria Lemos Nogueira, Ivanda de Souza Silva Tudesco, José Carlos F. Galduróz, Luciene Covolan, Orlando F. A. Bueno

## Abstract

The dopaminergic system is implicated in several cognitive processes including memory, attention and executive functions. This study was a double-blind, placebo-randomized trial designed to investigate the effect of dopamine D_2_ receptor blockade on episodic and working memory and particularly the relationship between executive functions, working memory capacity and long-term memory (LTM). Subjects ingested a single oral dose (4 mg) of haloperidol, a dopamine D_2_ receptor antagonist or placebo. Multiple linear regression using generalized linear models and a generalized estimating equation were used for statistical analyses. The results demonstrated that haloperidol impaired episodic memory (free recall of words and prose recall), working memory capacity-WMC (operation span task-OSPAN) and highly demanding executive functions (random number generation - RNG). In addition, it demonstrated that despite the large impairment in the RNG task performance in the haloperidol group, it did not affect episodic memory. The OSPAN task is predictive of episodic memory impairment, suggesting that memory impairments produced by haloperidol could be due in part to the impairment of WMC. As WMC partly relies on the appropriate functioning of the medial temporal lobe, probably the haloperidol-induced impairment on episodic memory through the decrease in the performance of WMC may depend on the activation of this area of the brain. The present study is relevant because it provides data on dopaminergic modulation of memory systems; suggesting that the major cause of deficits in episodic memory may be due to hippocampal function and WMC impairments, the latter more specifically with regard to controlled search and binding.

## INTRODUCTION

Episodic memories are formed by connecting information in a given time and space. These unrelated pieces of information bind together to form a representation of an integrated scene or event that can be consciously recalled or recognized [1]. In contrast, working memory (WM) is a limited-capacity system that simultaneously stores and manipulates short-term information. This system is important for attention, logic, reasoning, planning, strategy implementation and learning [2, 3].

Episodic memory and WM were traditionally considered both functionally and neuroanatomically dissociated [4, 5]. However, this view has been changing because of recent behavioral neuroimaging and electrophysiological findings [6-12]. Long-term memory (LTM) seems to contribute to WM, which in turn, contributes to episodic memory formation. This relationship is based on large-scale connection pathways between the pre-frontal cortex (PFC) and the temporal lobe, which are the brain regions intensively involved in WM and episodic memory, respectively [13].

The multi-component model proposed by Baddeley and Hitch [14] is an influential model of WM, that postulates two short-term storage loops: the phonological loop, which is capable of holding and processing verbal-based information, and the visuospatial sketchpad loop, which performs similar functions for visual information. Both loops are thought to be controlled by a central executive that allocates limited attentional resources. The executive functions include inhibition of prepotent responses, shifting mental sets, monitoring, updating task demands, goal maintenance and planning. In 2000, a new component was added to the model, the episodic buffer. This new version of the model proposed that the episodic buffer holdon-line information for short (although unspecified) periods and is responsible for storage capacity, prior considered an executive function [15]. The episodic buffer also appears to play an important role in sending new information to episodic memory and retrieving previously stored information from it[15].

According to Baddeley and Hitch [14], WM is a limited-capacity “workspace”, where the amount of storage required and the rate at which other processes can be carried out is limited [16]. More specifically, working memory capacity (WMC) refers to the relative capacity of this system to actively retain relevant information for a sufficient duration in the face of interference or distraction [17]. Tasks designed to assess WMC combine information storage and processing. The operation span (OSPAN) task, for example, requires a subject to carry out arithmetical operations while reading and maintaining words in memory for a short time [18]. The number of sets (1 word + 1 math problem) increases progressively in each list and range from 2-6. Therefore, these types of tasks demand a storage capacity that exceeds that of the phonological loop and the visuospatial sketchpad during the distracting process of performing an unrelated task (the distractor task). The distractor task (i.e solving math) prevents the subjectusing memory strategies to increase the number of items recalled (i.e. the words). Therefore, the OSPAN task provides a measure of WM span and reflects the storage capacity of the episodic buffer [19].

New information can interact with other information, like semantic, syntactic, visual and other perceptual details and through a process of “binding” and become one single representation during processing in short-term memory. This process, together with the chunking phenomenon which works to integrate items such as words and numbers, can greatly increase individual storage capacity. It is thought that different types of binding occur in the episodic buffer [20].

Regarding LTM, the roles of the hippocampus and its associated medial temporal lobe structures in the formation of episodic memories are well-established [21, 22]. Dopaminergic modulation has been reported during episodic memory encoding in humans [23] and decreased dopamine D_2_ receptor activation in the hippocampus has been implicated in memory and learning impairments in both preclinical [24, 25] and clinical studies [26-28]; for a negative result see Reeves [29]. However, few studies have been designed to assess the effects of dopamine D_2_ receptor blockade on episodic memory in healthy humans. This paucity of research is in sharp contrast to the numerous studies of dopamine D_2_ receptor antagonists in pathological states, as they are widely used in the treatment of pathologies such as schizophrenia[30] and Alzheimer’s disease [28, 31]conditions in which cognitive deficits are already present.

Concerning working memory, studies in monkeys and humans [32-35] have implicated dopamine D_2_ receptors in visual and spatial working memory, as haloperidol has been shown to reduce selective attention [36-38]. Dopamine D_2_ receptors have also been implicated in attentional processing and working memory due to their presence in the PFC [32-35, 39] and striatum [40-42]. But, few studies have investigated the role of the dopamine D_2_ receptor in WMC. For example, Gibbs and D’Esposito suggested that dopamine D_2_ receptor stimulation improves WM performance in individuals with low WMC [20]. However, the involvement of dopamine D_2_ receptors in the theoretically-based mechanisms of WM remains unclear.

Haloperidol may impair episodic memory by reducing the attentional resources that are required for appropriate encoding; or via impairing working memory capacity or by acting directly on dopamine D_2_ receptor in the hippocampus, which is a pivotal structure for episodic memory formation. The present study aimed to investigate the role of dopamine D_2_ receptor in both the episodic and working memory components in order to explore any possible associations between these types of memory, more specifically if executive functions and/or WMC performance influence LTM performance.

## METHODS

This study was a randomized double-blind, parallel placebo-controlled trial with single oral doses of placebo or haloperidol administered in identical capsules.

A double-blind study was chosen to avoid any bias in the results and the placebo control was used to have a comparative parameter between normal cognitive and altered dopaminergic conditions.

### Participants

Forty healthy young male volunteers participated in this study. All participants were native Portuguese speakers, 18-35 years of age, non-smokers, with normal body mass index (18<x<25 kg/m^2^) according to World Health Organization guidelines. The volunteers were undergraduate or graduate medical students (with at least 12 years of schooling) from the Universidade Federal de São Paulo. One week prior to the experiment, participants underwent a clinical interview, completed a Basic social and familiar questionnaire to verify their familial histories of disease, a Psychiatric state questionnaire and a Physical state questionnaire. At this time, intelligence quotient level was evaluated by a Raven Progressive Matrices test and basal anxiety level by the State Trait Anxiety Inventory (STAI-Trait) task. The participants were matched according to age, basal level of anxiety, education (years of schooling), and intelligence quotient. An expert psychiatrist acquired anamneses and performed clinical examinations. Participants, who reported histories of current or previous neurological or psychiatric disorders, current drug abuse, alcoholism, smoking, sleep disorders, or the use of any prescription medication, including vitamins and antioxidants, were excluded. Females were not included due to the possible effects of haloperidol on the tuberoinfundibular dopaminergic pathway, which may induce side effects, such as hyperprolactinemia, mastalgia and menstrual irregularities. All participants signed consent forms after being informed about the design of the study and the possible side effects of the drug. The Research Ethics Committee of the Universidade Federal de São Paulo approved the study (CEP N^0^. 0395/03).

### Treatment

Forty volunteers ingested a single oral dose of haloperidol (Haldol®, 4 mg, n=20) or placebo (lactose) (n=20). Haloperidol is a dopamine D_2_ receptor antagonist, a classical antipsychotic. This dosage was defined due to the high level of dopamine D_2_ receptor occupancy (approximately 73% after a single oral dose) [43-46], its efficacy in inducing cognitive effects, and its low incidence of extrapyramidal side effects[47, 48].

### Episodic memory tasks

Free recall of words task [49, 50] ten lists containing 15 unrelated concrete words from the Portuguese language were presented on a computer screen. Subjects read aloud the 15 sequentially presented words. After it, subjects recalled them as many as they could immediately (immediate recall) and 2 minutes later (delayed recall) in any order. Normally, subjects remember more of the words from the beginning (primacy effect) or the end of a word list (recency effect), producing a “U” shaped curve in a graph, known as a serial position curve [49-51]. The serial position curve refers to the graph relating the probability of recall with the position of the experimenter’s list. The serial position curve was analyzed in blocks of three words each, which resulted in 5 positions. The primacy effect results from the sub-vocal rehearsal of the initial words [49, 52] and the recency effect results from the temporary maintenance of information in the phonological loop (short-term memory). Thus, the recency effect is substantially reduced in the delayed recall condition [53].

The order of the lists were presented in 3 different ways to both groups (experimental and control) but the order of words in each list was the same. It is necessary to avoid that words previously studied in a list serve as a memory cues to the next list.The score was defined as the number of words correctly recalled for each block of 3 words position (block 1 to 5). The maximum possible score was 30 [the number of words contained in each block (3) vs. number of lists (10)].

Word lists are composed of independent items that do not have a meaning as a whole. However, the words are bound to the position they are in the list.

Prose recall task subjects were required to recall 14 sentences of a short prose passage immediately (immediate recall) or 30 minutes after (delayed recall) listening to the passage. Scores were obtained by awarding 1 point for each correctly recalled sentence or zero if containing any error or were not remembered[54, 55]. Prose recall allows episodic memory to be evaluatedwithin a defined context.

### Working memory and executive function tasks

Random number generation (RNG) task participants were required to generate out loud random sequences of numbers (1-10) within a given time interval (1/1 s) until 100 responses. Meanwhile the experimenter noted the numbers that had been spoken in a paper. They had to avoid talking numbers in order, what is difficult because in childhood in the learning of number, they were stored sequentially (1, 2, 3…). To accomplish this task, the participant handles the information in real time, inhibits habitual or stereotyped responses, generates new responses, and monitors and changes response-production strategies. The RNG task is a brief and efficient measure of executive functions (3) and is a clinically useful tool for assessing frontal lobe disturbances [56, 57].

The experimenterinsertedthe numbers in a software that evaluates the degree of randomness. It was scored using the Evans Index [58]. This index provides a sensitive measure of randomness (reflecting the disproportion with which any number follows any other number).So is, therefore, a measure of sequential response bias. The score may range between 0-1. The subject that does not randomize anything, i.e. repeats the same value 100 times, would have an index of 1. The higher score, the worse the performance.

*Operation span task (OSPAN) [18]* this task required the participants to verify arithmetical operations and memorize non related words. The number of arithmetic-word sequences may range from 2 to 6 and sequences of different length are presented in a random order to make it difficult for the subject to create memory strategies. The arithmetical operations alternate with the word presentations. The end of each set was followed by a question mark that indicated the moment at which the participant must decide if the operation were right or wrong and then read the subsequent word aloud, for example: [(4×5) + 5 = 31? Right or Wrong? Horse]. After each group of sets (2-6), participants were required to report all words sequentially. If they did not remember some words they should have said “Zum” instead. As the number of items in the operation-word sets increase, participants become increasingly unable to remember the words. The task evaluates how many items can be actively maintained in memory while attention is focused on another activity, i.e. solving mathematical operations. The OSPAN task measures working memory capacity, as it requires simultaneous processing and storing of information for a brief period [14, 59-61]. There were three different sequences of each group of set presented and randomly distributed among the volunteers (but the order of words in each set was the same). This is necessary to avoid that a sequence of set of words previously studied serve as a memory cues.

The task took about 10 min to be applied excluding the time for training for the subject to understand it. It could vary depending on the subject. Those who had ingested the drug were usually slower to perform the task. Scores is based on how many of the words (maximum 60) were perfectly recalled. 100% accuracy is represented by the value 1, and as the subject fails to recall the words, the value drops to a minimum value of 0. The mathematical operations accuracy was calculated but not considered in the final score. It served as a distracting task and to ensure that the participant engaged in the processing task once results were computed only if the participant solved at least 80% of the arithmetic operations.

### Procedure

Participants were asked to avoid consuming alcohol, energy drinks, chocolate, coffee or any other psychoactive substance for 24 hours prior to the experiment. They were instructed to have at least 7 hours sleep the previous night.

The experiment began at 7:30 a.m. Subjects had not eaten for 12 hours prior to the beginning of the experiment, and were given a controlled breakfast that was free of tryptophan, tyrosine, and caffeine 15 minutes after capsule ingestion. The participants took part ina brief training session to confirm they understood the instructions for each of the tasks. Neuropsychological tests were initiated 3 hours after the capsules were ingested; by this time, the peak haloperidol plasma level of 4 mg had been reached (the half-life of haloperidol is 26.5 ± 13.5 hours) [48].

During this three-hour interval, participants remained on site, either reading or watching TV. WM tasks were conducted after the free recall of words task.

### Statistical analyses

Multiple linear regression using generalized linear models (GLM) and the generalized estimating equation (GEE) were used to explore the main effects of the treatment on task performances. Distributions and link functions were selected by goodness of fit using Akaike information criterion (AIC) in the GLM and quasi-likelihood under the independence criterion (QIC) in the GEE.The analyses of free recall of words data, considered blocks 1 through 4 with block 5 being discarded (short-term memory). To evaluate the haloperidol treatment and the retention interval effects (immediate and delayed) on free recall of words and prose recall, GEE (with gamma distribution with log link function) were performed. Multivariate analyses using RNG or OSPAN as the covariate verified whether haloperidol and retention effects on free recall of words were independent of an OSPAN and or RNG task performance effect. The significance level adopted was p<0.05.

## RESULTS

The placebo and haloperidol groups had similar and normal body mass indices [F_(1,38)_= 0.75; p=0.39; Cohen’s *d*=0.28], similar mean ages [F_(1,38)_= 0.20; p=0.66; Cohen’s *d*=0.23], and the same basal levels of anxiety (STAI-trait scores) one week before the experiment [F_(1,38)_= 0.63; p=0.43; Cohen’s *d*=0.19] and on the day of the experiment [F_(1,38)_= 0.20; p=0.60; Cohen’s *d*=0.14] (Table 1).

**Table 1.**
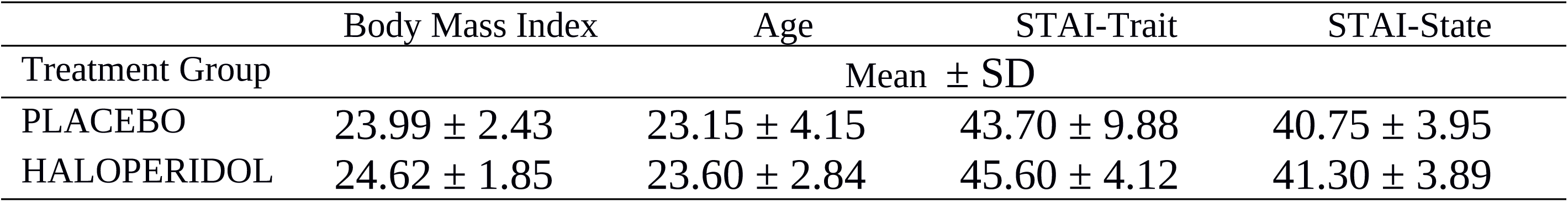
Demographic data and anxiety level of subjects.

|  | Body Mass Index | Age | STAI-Trait | STAI-State |
| --- | --- | --- | --- | --- |
| Treatment Group | Mean ± SD | | | |
| PLACEBO | 23.99 ± 2.43 | 23.15 ± 4.15 | 43.70 ± 9.88 | 40.75 ± 3.95 |
| HALOPERIDOL | 24.62 ± 1.85 | 23.60 ± 2.84 | 45.60 ± 4.12 | 41.30 ± 3.89 |

### Episodic memory tasks

#### Free recall of words

Significant effects of haloperidol were observed on the task (p<0.0001) in both the immediate and delayed (p<0.0001) recall tasks. The haloperidol group presented impairments in free recall of words when compared to the placebo group (Cohen’s d=0.93). Furthermore, performance was worse in delayed recall than in immediate recall in both groups (Cohen’s d= 0.76; Figs. 1, 2).

**Figure 1.**
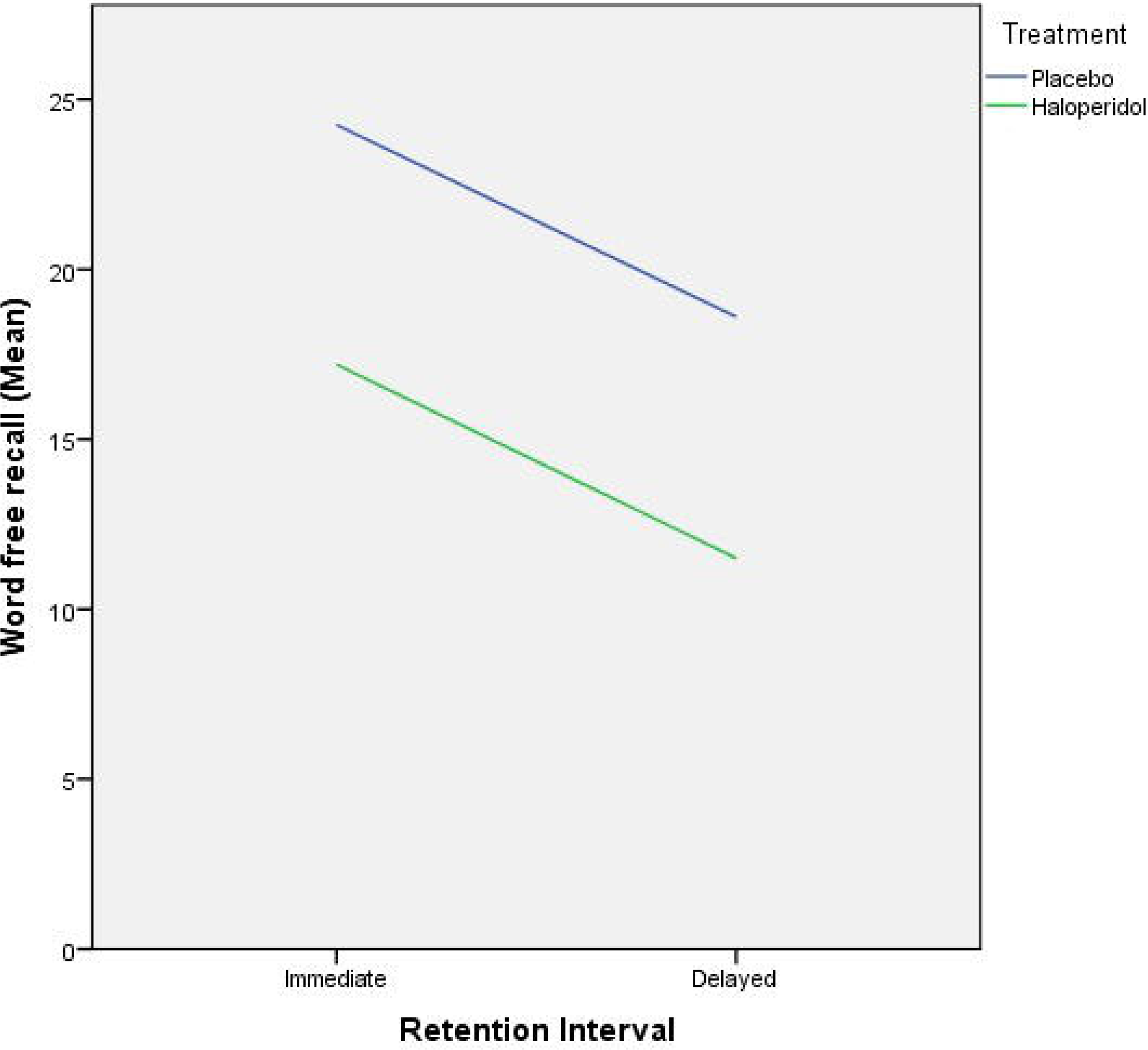
Free recall of words (block 5 excluded).

**Figure 2.**
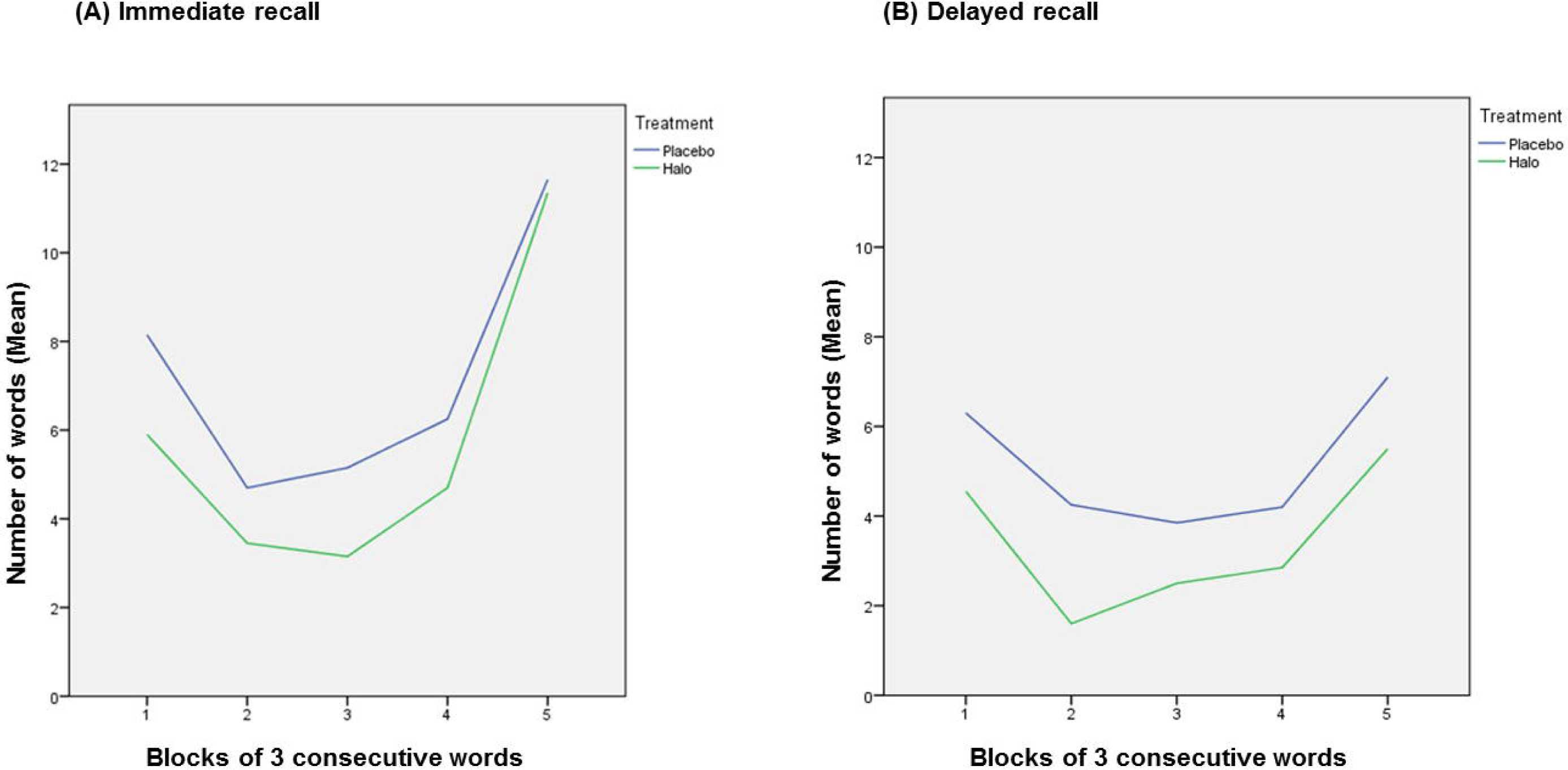
Serial position curve: A. Immediate recall, B. Delayed recall.

#### Prose Recall

Impairment on prose recall was observed in the haloperidol group when compared to the placebo group (p=0.012) and a significant effect of retention interval (immediate and delayed, p<0.0001) in prose recall score levels was also found. The haloperidol presented lower mean values (Cohen’s d=0.64) what means worse performance than the placebo group. Lower mean values were also observed in delayed prose recall than immediate prose recall in both groups (Cohen’s d=0.57; Fig. 3).

**Figure 3.**
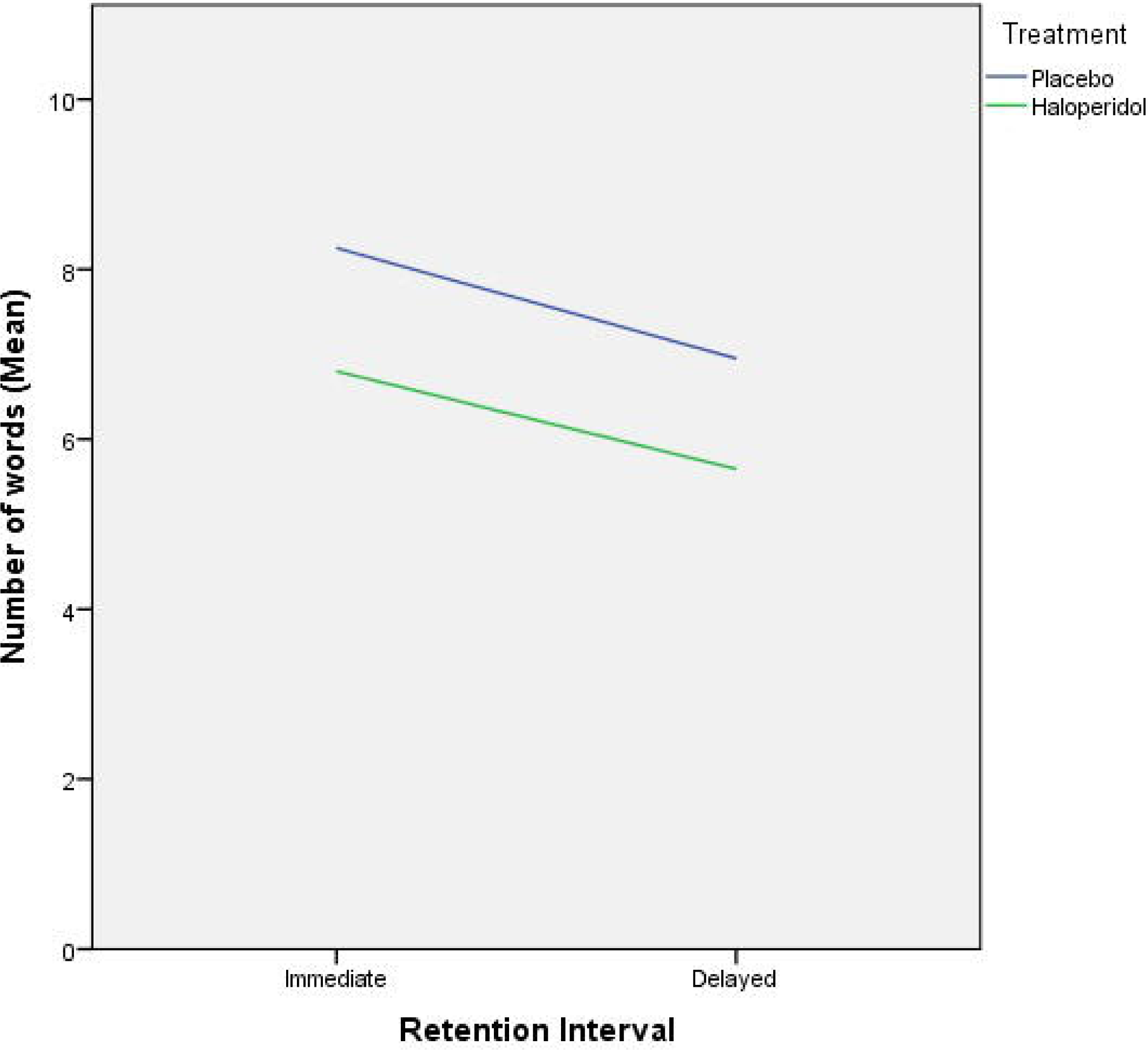
Prose recall performance.

### Working memory tasks

A GLM (with normal distribution with identity link function) revealed a significant effect of haloperidol on the OSPAN task (Walt=19.601, DF=1, p<0.0001). The haloperidol group presented lower mean values which represent worse performance than placebo group (Cohen’s d=1.13, 95% CI: 0.63 - 1.63; Fig. 4a). An effect of haloperidol on the RNG task was also found (Walt=59.3, DF=1, p<0.0001), as the haloperidol group showed the highest RNG values which represent worst performance (Cohen’s d=1.54, 95% CI: 1.13 - 1.92; Fig.4b). A correlation was found between RNG and OSPAN [Pearson correlation= -0.312; Sig. (2-tailed)= 0.050; Fig. 5]. Since working memory capacity is theoretically under the control of executive functions (15), the RNG task was calculatedas a predictor of the OSPAN task. A significant result (Walt=4.314, DF=1, p<0.0001, p = 0.038) demonstrated that RNG task performance can interfere with OSPAN task performance, so both processes are related.

**Figure 4.**
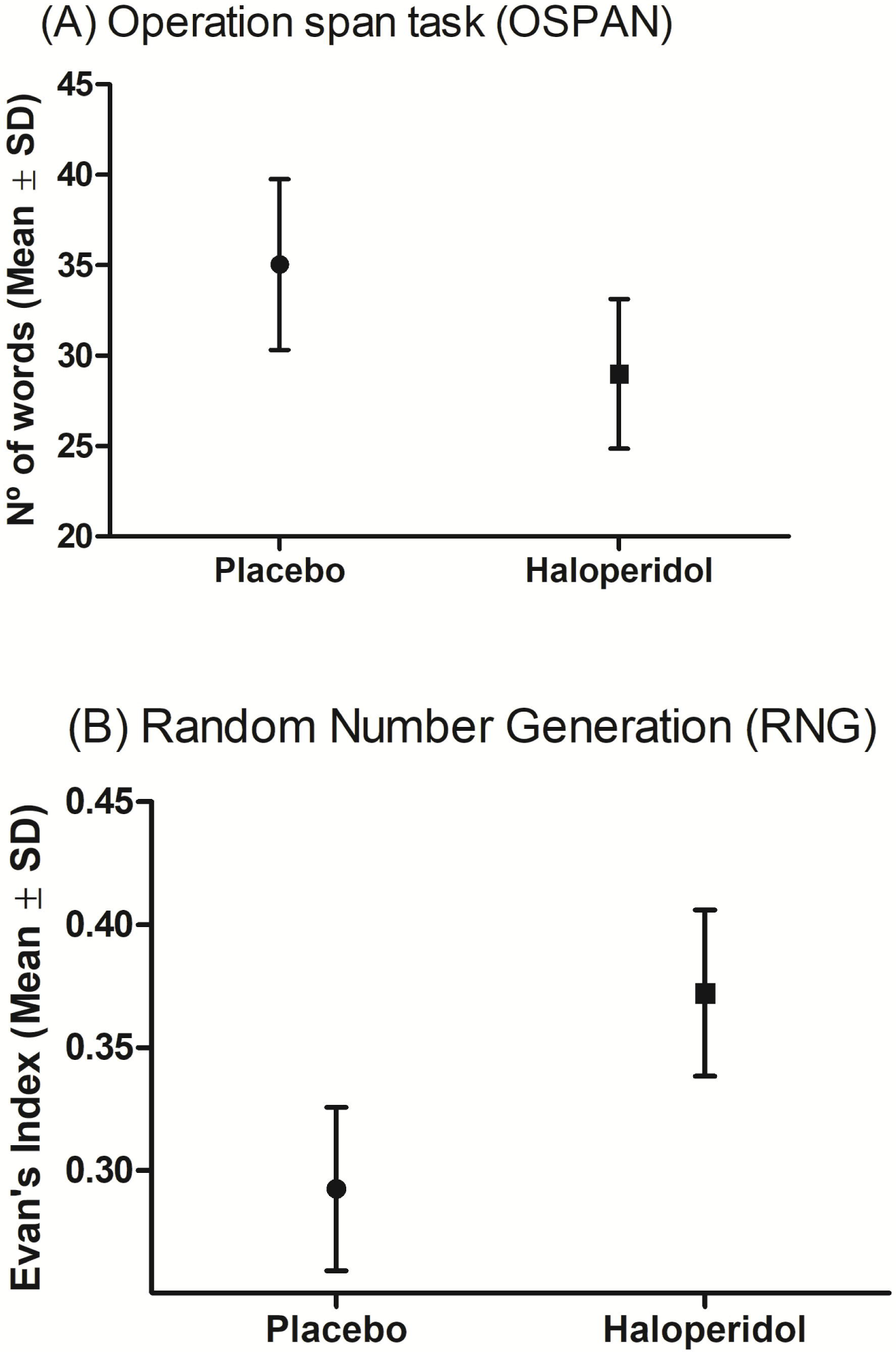
Haloperidol effect compared to placebo group on OSPAN and RNG (p<0.05).

**Figure 5.**
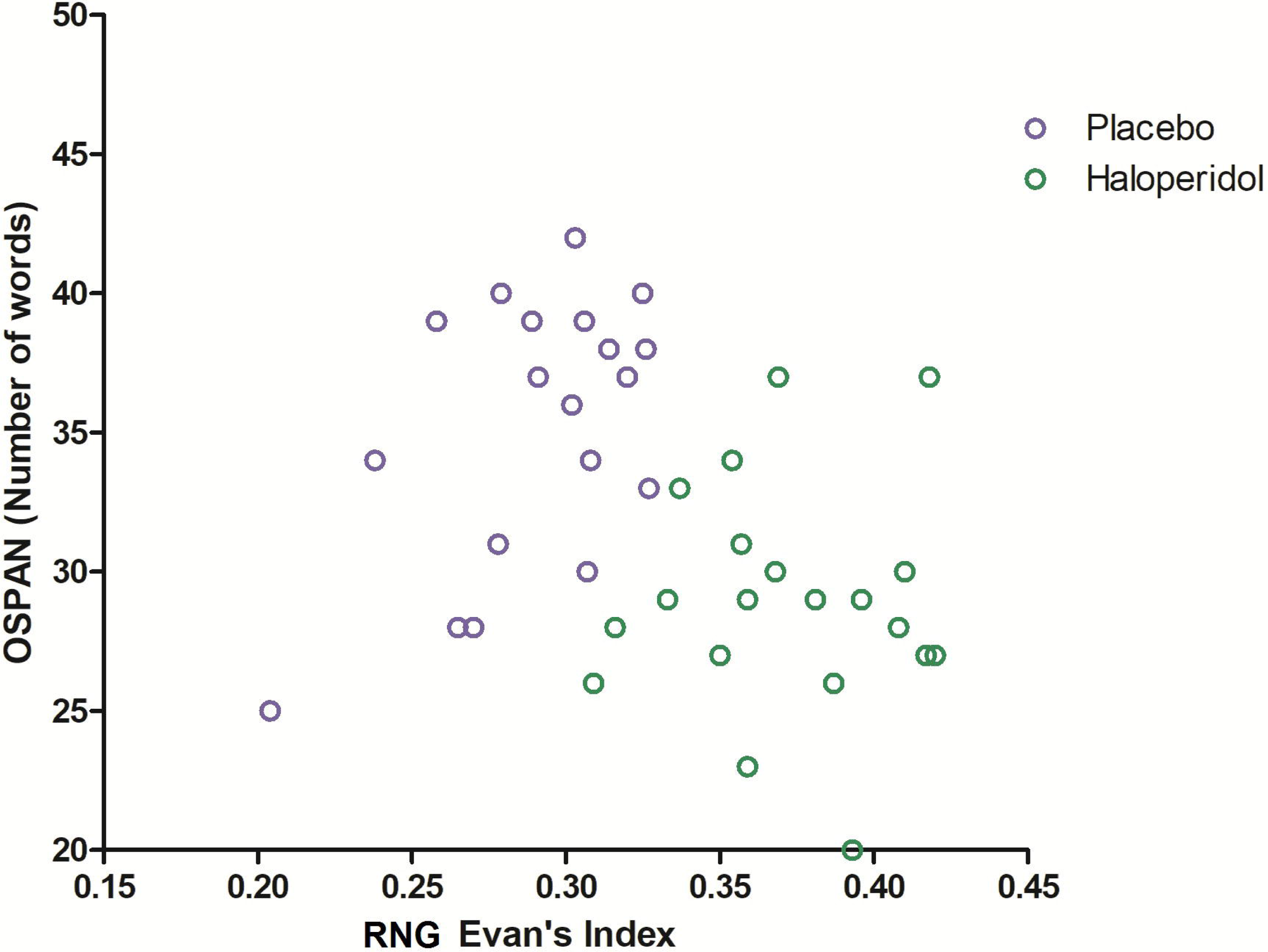
Correlation between OSPAN and RNG task.Placebo and haloperidol distribution.

Other multivariate analyses were performed to determine whether the effect of haloperidol on prose recall was independent of OSPAN or RNGperformance, in both immediate and delayed conditions. The results suggested a drug effect independent of both OSPAN and RNG (no power of test) performance on prose recall (p=0.081).

Multivariate analyses demonstrated that the effect of haloperidol on free recall of words for both the immediate and delayed conditions was independent of any RNG effect. The analyses revealed that OSPAN performance is predictive of free recall of words performance, whether in immediate recall or in delayed recall tasks. Therefore, WMC is relevant to the performance of episodic memory.

## DISCUSSION

Haloperidol impaired the episodic memory tasks (free recall of words and prose recall) with a retention interval effect being observed in both tasks.

Free recall of words taskimpairment was observed in all blocks of the serial position curve, with the exception of the last block. This block is subject to the recency effect and is usually interpreted as reflecting short-term memory processes [51, 62].In prose recall impairments were observed in both immediate and delayed recall.

Other agents, such as benzodiazepines and anticholinergic drugs, have also been shown to impair episodic memory, although the pharmacological mechanisms of action are entirely different [63, 64]. Given that the hippocampus and parahippocampal gyrus contain dopamine D_2_ receptors [65] and are brain structures critically related to episodic memory formation [22], it is likely that haloperidol-induced episodic memory impairment is caused by the direct action of the drug on these structures [65-67]. This hypothesis is in accordance with previously reported correlations between memory impairment and decreased dopamine D_2_ receptor binding in the hippocampus and parahippocampal gyrus in Alzheimer’s patients with severe amnesia [26-28]. The present study reported that an acute blockade of dopamine D_2_ receptor by haloperidol administration couldimpair episodic memory in healthy young subjects by affecting hippocampus and parahippocampal gyrustoo.

In addition, haloperidol induced a deficit in working memory performance as demonstrated by the impairment observed in the RNG and OSPAN tasks, which assess executive functions and WMC, respectively.

Haloperidol ingestion caused a large impairment in the RNG task, which is in line with a previous study that demonstrated a relationship among dopamine D_2_ occupancy (using raclopride), frontal metabolism and working memory. Jahanshahi and colleagues stated that the RNG task requires inhibition of habitual counting, i.e., the inhibition of prepotent responses acquired by previous overlearning[57, 68, 69]. Therefore, the present RNG results suggest a decrease in the ability of the individual to inhibit prepotent or irrelevant information during manipulation due to haloperidol administration.Inhibition is known to depend on PFC activity, particularly the left dorsolateral PFC [70]. However, the regression analyses in the present study showed no relationship between haloperidol impairment of RNG performance and haloperidol-induced episodic memory impairment. Inhibition of irrelevant information is, therefore, not by itself a predictor of episodic memory performance (free recall of words/prose recall).

The analysis pointed a positive correlation between RNG and OSPAN.The multivariate analysis with RNG as predictor of OSPAN indicatedthat the central executive contributes to working memory capacity performance. Loading and maintenance in working memory capacity require the inhibition of irrelevant information to keep relevant items in this temporary buffer, which is accomplished by PFC activity. Our results are consonant with prior studies that obtained a positive correlation between executive attention (primarily measured through inhibition) and WMC [71, 72].

The present study found that haloperidol decreased working memory capacity, as shown by performance in the OSPAN task.OSPAN is a working memory capacity test, as it indexes the amount of retained items in working memory in the face of distracting information. According to Baddeley’s framework, OSPAN task corresponds to the episodic buffer which holds integrated episodes or chunks in a multidimensional code. By doing so, it acts asstorage system which components of working memory, perception and episodic memory interact among themselves [15].

Regression analyses demonstrated that impaired OSPAN task performance (i.e., impairments of WMC) caused by dopamine D_2_ receptor blockade affected episodic memory performance (specifically the free recall of words task). This result in particular demonstrated that in the haloperidol group the relationship between WMC and episodic memory was not as strong as in the placebo group.WMC was, therefore, less predictive of episodic memory in the haloperidol group.

Deficits in episodic memory and WMC tasks have been shown to be correlated in healthy elderly subjects [9], and the present study observed the same result with healthy young subjects. According to some authors, WMC is important in controlled search processes of LTM and necessary for retrieval [73-75]. This strategic control process is thought to rely on executive functions and attention, including setting up a retrieval plan, retrieval strategies, monitoring strategies and decision-making and generating appropriate cues to search [73, 76]. Once a searched item is found, it is bound to other items (binding processing). These processes occur mainly in the prefrontal area, which is considered to be the primary mediator of working memory. Previous studies have suggested that baseline dopamine availability in this area is related to WMC[20], therefore, an explanation for the present results could be that dopamine D_2_ receptor blockade impaired the ability of the individual to use appropriate cues to access required information, which in turn compromised the binding process, resulting in impaired episodic memory. The amnesic effect of haloperidol on free recall of words may have been due to a double effect: the direct action of haloperidol on LTM and its effects on WMC. This would mean that there was a direct effect of haloperidol in the hippocampus and/or an indirect effect in the prefrontal cortex.

However, multivariate analyses failed to demonstrate that the effect of haloperidol on prose recall is dependent on its effect in OSPAN/RNG. Anderson and Bower proposed that during free recall of words the retrieval of items occurs according to certain context information originally stored with each item in the list[77]. This contextual information includes elements of the environment during the presentation of list, subject mood and posture. In prose recall, on the other hand, the information constitutes a context by itself, strong enough to resist the WMC impairment caused by dopamine D_2_ receptor blockade, at least at the dosage used in the present study. In prose recall performance, haloperidol treatment directly affected LTM independently of its effect on working memory subcomponents (working memory capacity/central executive) favoring the hypothesis that its effect was on the medial temporal lobe. If this is so, an explanation for the diverse effects of haloperidol on free recall of words and prose recall, both episodic memory tasks, should be searched for.

The lack of relationship among haloperidol impairment of RNG performance and haloperidol-induced episodic memories impairments reinforces the binding hypothesis, whichclaims that the working memorysystemcontrols, maintains, and updates arbitrary bindings. The limit of WMC arises from interference between bindings. The WM binding functions attributed to the WMC are more likely to reflect the drug complex action. Unsworth and Engle [74] asserted that WMC reflects a complex span performance with two components: (1) a *Maintenance process* that depends on limited storage capacity, (2) *Controlled retrieval* from memory over longer time periods. Both inhibition and memory-based abilities are important for successful retrieval, the first element necessary to keep irrelevant information out of the buffer, and the second to bind together the relevant information. According to Wilhelm et al. [75] “items in a list are bound to the list position, objects are bound to their position in space, and concepts are bound to roles in propositional schemata”, finally binding different stimuli from different domains. Independent items compound the lists and have no consistent meaning as a whole.Working memory is thought to be necessary to bind unrelated items of list with external context.The authors argued that high WMC reflects the ability to establish robust binding in WM.

Haloperidol must have compromised the binding of incoming words with context during the free recall of words in which each item needs to be encoded separately demanding more from the WMC than in the prose recall once during the binding of elements in prose recall they already have an internal context.What seems to be curious is that Wilhelm et al. claimed that high WMC reflects the ability to establish robust binding in WM [75] while Gibbs & D’Esposito [20] suggested that dopamine D_2_ receptor stimulation improves WM performance in individuals with low WMC. From our results dopamine blockade compromised more the lists (“weak binding?) than prose recall, considered more robust binding. So, could we suppose that dopamine D_2_ receptor would have a more important role in less robust binding? It is a matter to be solved in the future.

In summary, haloperidol compromised episodic memory, executive functions and working memorycapacity suggesting that these functions depend on dopamine D_2_receptors. In addition, WMC’s impairment contributed to worse performance of episodic memory, particularly in the task of free recall of words.

The study also reflects on the possibility that the drug has particularly affected the processes of inhibition, controlled search of memory cues and binding. In addition, the results of the study pointed out that the impairment of the WMC, important for controlled search and binding, may have been together with hippocampal function impairments the major cause of deficits in episodic memory.

Further studies with other tasks evaluating the same WMC domain (i.e., such as the N-back task, the running-memory task, and the memory-updating task) and other domains that provide answers about WM-LTM relationship as well as other drugs implied in related neurotransmitter systems are necessary to disentangle this issue.

## Acknowledgements

We would like to thank all the participants for their contributions to this study.

## Funding Acknowledgements

This work was supported by the Associação Fundo de Incentivo à Pesquisa (AFIP) [grant number: 130202] and Conselho Nacional de Desenvolvimento Científico e Tecnológico (CNPq) [grant number: 141383/2010-0], CNPq Grant level 1A.

## REFERENCES

1. Eichenbaum, H., Hippocampus: Cognitive processes and neural representations that underlie declarative memory. Neuron, 2004. 44: p. 109–120.

2. Baddeley, A., Working memory. Science, 1992. 255: p. 556–559

3. Baddeley, A. and S. Della Sala, eds. Working memory and executive control. The Prefrontal Cortex, ed. R.T. Roberts AC, Weisenkrantz L 1998, Oxford University Press: Oxford

4. Axmacher, N., et al., Sustained neural activity patterns during working memory in the human medial temporal lobe. The Journal of Neuroscience, 2007. 27: p. 7807–7816.

5. Campo, P., et al., Is medial temporal lobe activation specific for encoding long-term memory? Neuroimage, 2005. 25: p. 34–42.

6. Axmacher, N., et al., Interactions between medial temporal lobe, prefrontal cortex, and inferior temporal regions during visual working memory: a combined intracranial EEG and functional magnetic resonance study. The Journal of Neuroscience, 2008. 28: p. 7304–7312.

7. Axmacher, N., et al., Interaction of working memory and long-term memory in the medial temporal lobe. Cerebral Cortex, 2008. 18: p. 2868–2878.

8. Bueno, O., ed. Studying memory: from the frontal to the temporal lobe and vice-versa. Learning and Memory Development and Intellectual Disabilities, ed. L. Ecklund and A. Nyman. 2010, Nova Science: Hauppage. 225–238.

9. De Luccia, G., O. Bueno, and R. Santos, Recordação livre de palavras e memória operacional em idosos. Distúrbios da Comunicação, 2005. 17: p. 347–359.

10. Ranganath, C., M. Cohen, and C. Brozinsky, Working memory maintenance contributes to long-term memory formation: neural and behavioral evidence. Journal of Cognitive Neuroscience, 2005. 17: p. 994–1010.

11. Wagner, D., et al., Material-specific lateralization of working memory in the medial temporal lobe. Neuropsychologia, 2009. 47: p. 112–122.

12. Weiss, S. and P. Rappelsberger, Long-range EEG synchronization during word encoding correlates with successful memory performance. Cognitive Brain Research, 2000. 9: p. 299–312

13. Goldman-Rakic, P., L. Selemon, and M. Schwartz, Dual pathways connecting the dorsolateral prefrontal cortex with the hippocampal formation and parahippocampal cortex in the rhesus monkey. Neuroscience, 1984. 2: p. 719–743.

14. Baddeley, A. and G. Hitch, Working memory, in The Psychology of Learning and Motivation, B. GA, Editor. 1974, Academic Press: New York.

15. Baddeley, A., The episodic buffer: a new component of working memory? Trends in Cognitive Sciences, 2000. 4: p. 417–423.

16. Logie, R. and N. Cowan, Perspectives on working memory. Memory and Cognition, 2015. 43: p. 315–324

17. Engle, R., et al., Working memory, short-term memory and general fluid intelligence: a latent-variable approach. Journal of Experimental Psychology: General, 1999. 128: p. 309–331.

18. Turner, M. and R. Engle, Is working memory capacity task dependent? Journal of Memory and Language, 1989. 28: p. 127–154.

19. Baddeley, A., Working memory, thought and action. 2007, Oxford: Oxford University Press.

20. Gibbs, S. and M. D’ Esposito, Individual capacity differences predict working memory performance and prefrontal activity following dopamine receptor stimulation Cognitive Affective, & Behavioral Neuroscience, 2005. 5(2): p. 212–221.

21. Bertolucci, P., et al., Permanent global amnesia: case report. Clinical and Investigative Medicine, 2004. 27: p. 101–106.

22. Squire, L., Memory systems of the brain: A brief history and current perspective. Neurobiology of Learning and Memory, 2004. 82: p. 171–177.

23. Schott, B., et al., The dopaminergic midbrain participates in human episodic memory formation: evidence from genetic imaging Journal of Neuroscience, 2006. 26: p. 1407–1417.

24. Fujishiro, H., et al., Dopamine D2 receptor plays a role in memory function: implications of dopamine-acetylcholine interaction in the ventral hippocampus. Psychopharmacology, 2005 182: p. 253–261.

25. Umegaki, H., et al., Involvement of dopamine D(2) receptors in complex maze learning and acetylcholine release in ventral hippocampus of rats. Neuroscience, 2001. 103: p. 27–33.

26. Joyce, J., et al., Dopamine D2 receptors in the hippocampus and amygdala in Alzheimer’s disease. Neuroscience Letters, 1993. 154: p. 171–174.

27. Joyce, J., A. Myers, and E. Gurevich, Dopamine D2 receptor bands in normal human temporal cortex are absent in Alzheimer’s disease. Brain Research, 1998. 784: p. 7–17.

28. Kemppainen, N., et al., Hippocampal dopamine D2 receptors correlate with memory functions in Alzheimer’s disease. European Journal of Neuroscience, 2003. 18: p. 149–154.

29. Reeves, S., et al., The dopaminergic basis of cognitive and motor performance in Alzheimer’s disease. Neurobiology of Disease, 2010. 37: p. 477–482

30. Suwa, H., et al., Attention disorders in schizophrenia. Psychiatry and Clinical Neurosciences, 2004. 58: p. 249–256.

31. Martorana, A. and G. Koch, Is dopamine involved in Alzheimer Disease? Frontiers in Aging Neuroscience, 2014. 6: p. 1–6.

32. Arnsten, A., et al., Dopamine D2 receptors mechanisms contribute to age-related cognitive decline: the effects of quinpirole on memory and motor performance in monkeys.. The Journal of Neuroscience, 1995. 15: p. 3429–3439.

33. Luciana, M. and P. Collins, Dopaminergic modulation of working memory for spatial but not object cues in normal volunteers. Journal of Cognitive Neuroscience, 1997. 9: p. 330–347.

34. Luciana, M., et al., Facilitation of working memory in humans by a D2 dopamine receptor agonist. Journal of Cognitive Neuroscience, 1992. 4: p. 58–67.

35. Mehta, M., et al., Systemic sulpiride in young adult volunteers simulates the profile of cognitive deficits in Parkinson’s disease. Psychopharmacology, 1999. 146: p. 162–74.

36. Ahveninen, J., et al., Suppression of transient 40-Hz auditory response by haloperidol suggests modulation of human selective attention by dopamine D2 receptors. Neuroscience Letters, 2000. 292: p. 29–32.

37. Coull, J., et al., Differential effects of clonodine, haloperidol, diazepam, and tryptophan depletion on focused attention and attentional search. Psychopharmacology, 1995. 121: p. 222–230.

38. Kähkönen, S., et al., Effects of haloperidol on selective attention. Neuropsychopharmacology, 2001. 25: p. 498–504.

39. Takahashi, H., et al., Differential contributions of prefrontal and hippocampal dopamine D1 and D2 receptors in human cognitive functions. J Neurosci, 2008. 28: p. 12032–12038.

40. Bäckman, L., et al., Age-related cognitive deficits mediated by changes in the striatal dopamine system. Am J Psychiatry, 2000. 157: p. 635–637.

41. Cervenka, S., et al., Associations between dopamine D2-receptor binding and cognitive performance indicate functional compartimentalization of the human striatum. Neuroimage, 2008. 40: p. 1287–1295.

42. Volkow, N., et al., Association Between Decline in Brain Dopamine Activity With Age and Cognitive and Motor Impairment in Healthy Individual. American Journal of Psychiatry, 1998. 155: p. 344–349.

43. Forsman, A. and R. Öhman, Pharmacokinetics studies on haloperidol in man. Current Therapeutical Research, Clinical and Experimental, 1976. 20: p. 319–336.

44. Kapur, S., et al., High levels of dopamine D2 receptor occupancy with low-dose haloperidol treatment: a PET study. The American Journal of Psychiatry, 1996. 153: pp. 948–950.

45. Nordström, A., L. Farde, and C. Halldin, Time course of D2-dopamine receptor occupancy examined by PET after single oral doses of haloperidol. Psychopharmacology, 1992. 106: p. 433–438.

46. Nordström, A., et al., Central D2-dopamine receptor occupancy in relation to antipsychotic drug effect: A double blind PET study of schizophrenic patients.. Biological Psychiatry, 1993. 33: p. 227–235.

47. Braude, W., T. Barnes, and S. Gore, Clinical Characteristics of Akathisia. A systematic investigation of acute psychiatric inpatients admissions. The British Journal of Psychiatry, 1983. 143: p. 139–150.

48. King, D., The effect of neuroleptics on cognitive and psychomotor function. British Journal of Psychiatry, 1990. 157: p. 799–811.

49. Capitani, E., et al., Recency, primacy and memory: reappraising and standardising the serial position curve. Cortex, 1992. 28: p. 315–342.

50. Glanzer, M. and A. Cunitz, Two storage mechanisms in free recall. Journal of Verbal Learning and Verbal Behavior, 1966. 5: p. 351–360.

51. Murdock, B., The serial position effect of the free recall. Journal of Experimental Psychology, 1962. 64: p. 482–483.

52. Rundus, D., Analysis of rehearsal processes in free recall. Journal of Experimental Psychology, 1971. 89: p. 63–77.

53. Baddeley, A., ed. Human memory: Theory and Practice 1997, Psychology Press: Hove

54. Correa, D. and C. Gorenstein, Bateria de testes de memória. Parte 2. Critérios de elaboração e avaliação. Arquivos Brasileiros de Psicologia, 1988. 40: p. 42–53.

55. Correa, D. and C. Gorenstein, Bateria de testes de memória. Parte 1. Critérios de elaboração e avaliação. Arquivos Brasileiros de Psicologia, 1988. 38: p. 24–35.

56. Brugger, P., et al., Random number generation in dementia of the Alzheimer type: A test of frontal executive functions. Neuropsychologia, 1996. 34(97-103).

57. Jahanshahi, M., et al., The effects of transcranial magnetic stimulation over the dorsolateral prefrontal cortex on suppression of habitual counting during random number generation. Brain, 1998. 121: p. 1533–1544.

58. Evans, F., Monitoring attention deployment by random number generation: an index to measure subjective randomness. Bulletin of the Psychonomic Society, 1978. 2: p. 35–38.

59. Conway, A., et al., A latent variable analysis of working memory capacity, short-term memory capacity, processing speed, and general fluid intelligence. Intelligence, 2002. 30: p. 163–183.

60. Daneman, M. and P. Carpenter, Individual differences in WM and reading. Journal of Verbal Learning and Verbal Behavior, 1980. 19: p. 450–466.

61. Engle, R., ed. What is working memory capacity? The nature of remembering: Essays in honor of Robert G. Crowder, ed. H. Roediger, et al. 2001, American Psychological Association Press: Washington.

62. Brelsford, J., R. Freund Jr, and D. Rundus, Recency judgments in a short-term memory task. Psychonomic Science, 1967. 8: p. 247–248.

63. Curran, H., Benzodiazepines, memory and mood: a review Psychopharmacology, 1991. 105: p. 1–8.

64. Nogueira, A., et al., Effects of benzodiazepine on free recall of semantic related words.. Human Psychopharmacology Clinical and Experimental, 2006. 21: p. 1–10

65. Takahashi, H., et al., Memory and frontal lobe functions; possible relations with dopamine D2 receptors in the hippocampus. Neuroimage, 2007. 34: p. 1643–1649.

66. Goldsmith, S. and J. Joyce, Dopamine D2 receptor expression in hippocampus and parahippocampal cortex of rat, cat, and human in relation to tyrosine hydroxylase-immunoreactive fibers Hippocampus, 1994. 4: p. 354–373.

67. Ito, H., et al., Normal database of dopaminergic neurotransmission system in human brain measured by positron emission tomography. Neuroimage, 2008. 39: p. 555–565.

68. Jahanshahi, M. and G. Dirnberger, The left dorsolateral prefrontal cortex and random generation of responses: studies with transcranial magnetic stimulation. Neuropsychologia, 1999. 37: p. 181–190.

69. Jahanshahi, M., et al., Random number generation as an index of controlled processes. Neuropsychology, 2006. 20: p. 391–399.

70. Jahanshahi, M., et al., The role of the dorsolateral prefrontal cortex in random number generation: a study with positron emission tomography. Neuroimage, 2000. 12: p. 713–725.

71. Unsworth, N., On the division of working memory and long-term memory and their relation to intelligence: A latent variable approach. Acta Psychologica, 2010. 134: p. 16–28.

72. Unsworth, N. and G. Spillers, Working memory capacity: Attention control, secondary memory, or both? A direct test of the dual-component model. Journal of Memory and Language, 2010. 62: p. 392–406.

73. Unsworth, N., G. Brewer, and G. Spillers, Working memory capacity and retrieval from long-term memory: the role of controlled search. Memory and Cognition 2013. 41: p. 242–254.

74. Unsworth, N. and R. Engle, The Nature of Individual Differences in Working Memory Capacity: Active Maintenance in Primary Memory and Controlled Search From Secondary Memory. Psychological Review 2007. 114 (1): p. 104 –132.

75. Wilhelm, O., A. Hildebrandt, and K. Oberauer, What is working memory capacity, and how can we measure it? Frontiers in Psychology, 2013. 4: p. 1–22.

76. Nelson, T. and L. Narens, Metamemory: A theoretical framework and new findings, ed. T. Nelson and L. Narens. 1990, San Diego: San Diego Academic Press.

77. Anderson, J. and G. Bower, Recognition and retrieval processes in free recall. Psychological review, 1972. 79: p. 97–123.

